# A Complete Logical Approach to Resolve the Evolution and Dynamics of Mitochondrial Genome in Bilaterians

**DOI:** 10.1101/098764

**Authors:** Laurent Oxusoff, Pascal Préa, Yvan Perez

**Affiliations:** Laboratoire des Sciences de l’Information et des Systèmes UMR 7296, Aix-Marseille Université, Université de Toulon, CNRS, ENSAM, Marseille, France; Laboratoire d’Informatique Fondamentale de Marseille UMR 7269, Aix-Marseille Université, CNRS, Ecole Centrale Marseille, Technopole de Chateau-Gombert, Marseille, France; Institut Méditerranéen de Biodiversité et d’Ecologie marine et continentale UMR 7263, Aix-Marseille Université, Avignon Université, CNRS, IRD, Marseille, France

## Abstract

A new method of genomic maps analysis based on formal logic is described. The purpose of the method is to 1) use mitochondrial genomic organisation of current taxa as datasets 2) calculate mutational steps between all mitochondrial gene arrangements and 3) reconstruct phylogenetic relationships according to these calculated mutational steps within a dendrogram under the assumption of maximum parsimony. Unlike existing methods mainly based on the probabilistic approach, the main strength of this new approach is that it calculates all the exact tree solutions with completeness and provides logical consequences as very robust results. Moreover, the method infers all possible hypothetical ancestors and reconstructs character states for all internal nodes (ancestors) of the trees. We started by testing the method using the deuterostomes as a study case. Then, with sponges as an outgroup, we investigated the mutational network of mitochondrial genomes of 47 bilaterian phyla and emphasised the peculiar case of chaetognaths. This pilot work showed that the use of formal logic in a hypothetico-deductive background such as phylogeny (where experimental testing of hypotheses is impossible) is very promising to explore mitochondrial gene rearrangements in deuterostomes and should be applied to many other bilaterian clades.

**Author Summary:** Investigating how recombination might modify gene arrangements during the evolution of metazoans has become a routine part of mitochondrial genome analysis. In this paper, we present a new approach based on formal logic that provides optimal solutions in the genome rearrangement field. In particular, we improve the sorting by including all rearrangement events, *e.g*., transposition, inversion and reverse transposition. The problem we face with is to find the most parsimonious tree(s) explaining all the rearrangement events from a common ancestor to all the descendants of a given clade (hereinafter PHYLO problem). So far, a complete approach to find all the correct solutions of PHYLO is not available. Formal logic provides an elegant way to represent and solve such an NP-hard problem. It has the benefit of correctness, completeness and allows the understanding of the logical consequences (results true for all solutions found). First, one must define PHYLO (axiomatisation) with a set of logic formulas or constraints. Second, a model generator calculates all the models, each model being a solution of PHYLO. Several complete model generators are available but a recurring difficulty is the computation time when the data set increases. When the search of a solution takes exponential time, two computing strategies are conceivable: an incomplete but fast algorithm that does not provide the optimal solution (for example, use local improvements from an initial random solution) or a complete – and thus not efficient – algorithm on a smaller tractable dataset. While the large amount of genes found in the nuclear genome strongly limits our possibility to use of formal logic with any conventional computer, we show in our paper that, for bilaterian mtDNAs, all the correct solutions can be found in a reasonable time due to the small number of genes.

## Introduction

Unlike nuclear genomes, mitochondrial genomes (mtDNAs) are rather small and simply structured. In eukaryotes, they are considered relics of bacterial genomes and consist of circular DNA about 16 kb in size that, as a result of ancient intracellular symbiosis, have only retained a few well-characterized genes: 13 protein subunits (nad1-6, nad4L, cox1-3, cob and atp6/8), 2 rRNAs (rrnL, rrnS) and a maximum of 22 tRNAs [1]. Recently, the emergence of next-generation sequencing techniques has significantly increased the amount of mtDNAs available in public databases. The comparative analysis of this growing amount of data has helped to broaden our understanding of the evolution of mtDNAs in metazoans. Because it is assumed that nuclear genomes underwent similar evolutionary processes, it has been proposed that comparative analysis of mtDNAs could shed a new light on the mechanisms and selective forces driving whole-genome evolution in genomic data that are more tractable [2].

MtDNAs have also proven valuable for evolutionary studies in eukaryotes [1, 3–5]. Primary sequences from mitochondrial (mt) genes and mt control regions have not only yielded significant insights into the phylogenetic relationships of numerous taxa but have also been informative for population genetic investigations due to their maternal inheritance and high mutation rates. However, mutation rates vary considerably between species and several studies of metazoan relationships have suggested that they can be problematic to infer phylogenies [6].

Besides the primary sequence information, the organisation of the genome itself has also proven to be a reliable marker for phylogenetic inferences at many taxonomic levels for several reasons [4, 7]. First, the gene content is almost invariant in metazoans and provides a unique and universal dataset. Second, stable structural gene rearrangements are assumed to be rare because functional genomes must be maintained, which limits the level of homoplasy [8]. However, relying on mt gene order to make phylogenetic inferences has, at times, been disappointing because no evolutionary significant changes of gene order could be identified in some lineages [9]. There are several examples where mt gene orders were successfully used to support phylogenetic hypotheses, for instance crustaceans [10–12], echinoderms [13] and annelids [14, 15]. Nearly 80% of the rearrangements affect only tRNA genes. In the majority of these cases, only a single tRNA is affected [16]. The study of rearrangements of tRNA genes in the Hymenoptera suggests that the position of mt tRNA genes is selectively neutral [17]. Mt gene order can strongly differ within some lineages such as urochordates, molluscs, brachiopods, platyhelminths, bryozoans and nematodes [6]. The relationships of these shuffled genes phyla cannot be determined on the basis of gene order evidence and it should be noted that these phyla appear as long-branched leaves in sequence-based phylogenetic analyses [9, 18, 19], confirming that their molecular evolution rates are unusually fast. Mt gene rearrangements can be assigned to three main models: intrachromosomal recombination [20], Tandem Duplication followed by Random Loss of genes (TDRL model [3]) and a variant of the latter which consists of tandem duplication followed by non-random loss [21]. TDRL can easily describe local transposition but inversion and long-range transposition are more consistent with the intrachromosomal recombination model [20, 22]. TDRL events mostly occur across vertebrate lineages [22] and cannot explain gene inversions and long-range transpositions, which are common in invertebrate mtDNAs [24]. In protostomes, TDRLs represent about 10% of the rearrangement events [25]. Similar frequencies have been observed in the reconstructed rearrangements of metazoans. This may suggest that TDRL plays a marginal role (at least in invertebrates) and that recombination is an important component of the mtDNA rearrangement mechanism in metazoans [22]. There are four types of events in the intrachromosomal recombination model: inversion, transposition, reverse transposition (a transposition in which the re-inserted fragment is reversed) and gain/loss. Intrachromosomal recombination often involves the replication origins [23], but other hot spots of rearrangements have been described [26]. Consequentially, it is reasonable to consider intrachromosomal recombination events as elementary to the evolution of mtDNAs even though some transpositions may mechanistically result from TDLRs.

A variety of software tools implementing methods that automate the comparative analysis of gene order have been developed to infer phylogenies and genome evolution from mtDNAs [27, 28]. Most of these methods are based on the breakpoint number or inversion distance [29–38], and also integrate transposition events [39]. These methods are fast because they are based on approximations. Exhaustive exploration methods have rarely been exploited because they are NP-hard. The use of the web-based program CREx which considers transpositions, inversions, reverse transpositions, and TDRLs was recently proposed as one way to do so [25,40]. Alternative approaches based on character coding for parsimony and distance analyses from genomic rearrangements have been proposed to infer phylogeny [41, 42]. Finally, a cladistic approach to code gene order data allows the reconstitution of a possible hypothetical common ancestor genome at each node of the tree [43].

Here, we present the first logical study of mtDNAs organisation in bilaterians from pairs of mt gene orders. The goal of this new approach is to reveal the evolutionary history of the mt genomic rearrangements and exhibit ancestral gene orders (ground patterns). First, we used deuterostome mtDNAs as a study case. Second, we extended the analysis to bilaterians and emphasized the peculiar case of chaetognaths.

## Results and Discussion

Four useful definitions (3 to 6) towards the demonstration of the two properties below are given in Methods.

### Shared block property

#### Proposition

Let *A*_0_ and *A_k_* be two genomes such that the minimal distance d(*A*_0_, *A_k_*) is equal to *k*. There exists a path *C* = (*A*_0_, *A*_1_,…*A_k_*) between *A*_0_ and *A_k_* of minimal size (*i.e*., of *k* steps) and with no cut (see Definition 6). That is to say that in *C*, a block of genes present in two genomes *A_i_* and *A_j_* (possibly inverted) is always present (possibly inverted) in all the intermediate states between *A_i_* and *A_j_*.

#### Proof

Let *C* = (*A*_0_, *A*_1_,…*A_k_*) be a minimal path between *A*_0_ and *A_k_*. Let us suppose that in *C*, there is (at least) one cut, and that the last cut is between *A_i_* and *A*_*i*+1_ (the blocks shared between *A*_*i*+1_ and *A_k_* will not be cut in the path between *A*_*i*+1_ and *A_k_*). Let us call *G* = [*G*_1_ *G*_2_] the relevant block. *G* is cut (in *A_i_*) between *G*_1_ and *G*_2_: it exists *k′* in [*i*+2,…*k*] such that *G*_1_ and *G*_2_ are successive in *A_i_* and in *A*_*k′*_ but not in *A*_*i*+1_ nor in *A*_*k′*-1_.

We are in one of the following two cases:

##### Case 1

The mutation *μ_i_* between *A_i_* and *A*_*i*+1_ moves the block *G*_1_. So the block *G*_2_ does not move (otherwise, the mutation *μ_i_* is not a cut). Note that the case “*G*_2_ moves and not *G*_1_” is symmetrical. The mutation *μ_i_* moves the block *G*_1_ with a block *B* (possibly empty), neighbour of *G*_1_. The blocks *B* and *G*_1_ are successive in *A_i_* and *A*_*i*+1_.

We construct the path *C′* = (*A,′*_0_ = *A*_0_, *A′*_1_,…*A′*_*k*-1_, *A′*_*k*_ = *A_k_*) as follows:

1. For *j* from 0 to *i*: *A′*_*j*_ = *A_j_*
2. For *i* ≤ *j* < *k′*: the mutation *μ_j_* (in *C*) moves a block *E*=[*B G*_1_] containing the block *G*_1_ and not the block *G*_2_. The mutation *μ′_j_* (in *C′*) moves a block *E′*, obtained from *E* by deleting *G*_1_ (if *E′* is empty, it is equivalent to removing a step in the path).
3. For *j* ≥ *k′*: *μ′_j_* = *μ_j_* (others mutations are unchanged)

##### Case 2

The mutation *μ_i_* between *A_i_* and *A*_*i*+1_ does not move the block *G*_1_ nor the block *G*_2_. It moves a block *B* to the insertion point *x* located between *G*_1_ and *G*_2_.

We construct the path *C′* = (*A′*_0_ = *A*_0_, *A′*_1_,…*A′*_*k*-1_, *A′_k_* = *A_k_*) as follows:

1. For *j* from 0 to *i*: *A′_j_* = *A′_j_*
2. At Step *i*: the mutation *μ′_j_* moves (in *C′*) the block *B* to point *y* (instead of *x*), located at the “beginning” of *G*_1_ (*G*_1_ is located between *x* and *y*), with the same sign.
3. For *i* <*j* <*k′*:

a. If the mutation *μ_j_* moves (in *C*) a block *E* containing *G*_1_ and not *G*_2_, then, in *C′*, the mutation *μ′_j_* moves a block *E′*, obtained from *E* by removing *G*_1_.
b. If the mutation *μ_j_* moves (in *C*) a block *E* containing *G*_2_ (with or without *G*_1_), then, in *C′*, the mutation *μ′_j_* moves a block *E′*, obtained from *E* by replacing *G*_2_ by [*G*_1_ *G*_2_].
4. For *j* ≥ *k′*, *μ′_j_* = *μ_j_* (other mutations are unchanged).

In both cases, we obtained a path *C′* from *A*_0_ to *A*_k_, not longer than *C*, and containing less cuts. We get a path *C*\* from *A*_0_ to *A*_k_, not longer than *C* and containing no cuts by iterating the process. Starting from a minimum length path proves the result.

###### QED

The shared block property was used in the program genome_comparison.c as a heuristic test to decrease the calculation time. At each step of the computation of the shortest path between a genome *A* and a genome *B*, the blocks that are shared between the current genome *X* and the genome *B* are calculated first; the only mutations that are considered to be potentially applied to the current genome *X* are those that do not intersect the blocks shared between *X* and *B*. This property is reinforced by the following conjecture: let *A* and *B* be two genomes, *all* the shortest paths between *A* and *B* have no cut.

### Lower bound for minimal distance property

The property of the lower bound for minimal distance property defines a lower bound for the minimal distance between two genomes *A* and *B*, below which there is no path solution.

#### Proposition

Let *A* and *B* be two genomes at minimal distance *k* one from the other. We have 3 × *k* ≥ *nb_breakpoints*(*A*, *B*).

Examples:

If *nb_breakpoints*(*A*, *B*) = 4, 5, or 6, then d(*A*, *B*) ≥ 2

If *nb_breakpoints*(*A*, *B*) = 7, 8, or 9, then d(*A*, *B*) ≥ 3

If *nb_breakpoints*(*A*, *B*) = 10, 11, or 12, then d(*A*, *B*) ≥ 4

If *nb_breakpoints*(*A*, *B*) = 13, 14, or 15, then d(*A*, *B*) ≥ 5

#### Proof

Let *C* = (*A*_0_, *A*_1_,…,*A_i_*, *A*_*i*+1_,…,*A_k_*) be an arbitrary path (of length *k*) between genomes *A*_0_ and *A_k_*, and let μ_*i*_ be the mutation between *A_i_* and *A*_*i*+1_. If μ_*i*_ is a transposition or a reverse transposition, then only three intervals between two genes of *A_i_* are modified by μ_*i*_. Thus at most three breakpoints of *A_i_* are not breakpoints of *A*_*i*+1_. If μ_*i*_ is an inversion, then only two intervals are modified by μ_*i*_, and thus at most two breakpoints of *A_i_* are not breakpoints of *A*_*i*+1_. Thus, the number of breakpoints in *A*_0_ is at most 3*k*.

##### QED

This property was used in the program genome_comparison.c as a heuristic test to decrease the calculation time on the basis of the following contrapositive: Let *A* and *B* two distinct genomes. Let *k* > 0. If *nb_breakpoints*(*A*, *B*) > 3 × *k*, then there is no path between *A* and *B* of length *k*. Indeed, for each current genome *X* explored in the state 0 of the automaton (progression state), the function HEURISTIC_TEST returns NO if *nb_breakpoints*(*X*, *B*) > 3 × (number of steps remaining between *X* and *B*). In this case, the algorithm does not need to explore the search subtree from the current path, as it never leads to *B* with the remaining number of steps.

### Exact minimal distances and saturation threshold

Using the two previous properties as heuristics, it was possible to calculate the minimal distances between all pairs of genomes present in the taxonomic dataset (69 taxa, Table 1) considering transposition, inversion and reverse transposition. In contrast to previously published methods based on breakpoint number or inversion distance, the distance matrix obtained here is exact (S1 appendix).

**Table 1.** Species, systematic position, and accession number of mitochondrial genome used for gene order comparisons.

| Species | Taxonomy | Accession no. |
| --- | --- | --- |
| *Tethya actinia* | Porifera | AY_320033 |
| *Sipunculus nudus* | Sipuncula | FJ_422961 |
| *Urechis caupo* | Echiura | NC_006379 |
| *Platynereis dumerilii* | Annelida/Polychaeta | NC_000931 |
| *Phoronis architecta* (syn *psammophila*) | Phoronida | AY368231 |
| *Terebratalia transversa* | Brachiopoda | NC_003086 |
| *Terebratulina retusa* | Brachiopoda | NC_000941 |
| *Laqueus rubellus* | Brachiopoda | NC_002322 |
| *Lingula anatina* | Brachiopoda | AB178773 |
| *Gyrodactylus derjavenoides* | Platyhelminthes/Trematoda | NC_010976 |
| *Schistosoma mansoni* | Platyhelminthes/Trematoda | NC_002545 |
| *Katharina tunicata* | Mollusca/Polyplacophora | NC_001636 |
| *Biomphalaria glabrata* | Mollusca/Gastropoda | NC_005439 |
| *Cepaea nemoralis* | Mollusca/Gastropoda | NC_001816 |
| *Albinaria soerulea* | Mollusca/Gastropoda | NC_001761 |
| *Nautilus macromphalus* | Mollusca/Cephalopoda | NC_007980 |
| *Loligo bleekeri* | Mollusca/Cephalopoda | NC_002507 |
| *Siphonodentalium lobatum* | Mollusca/Scaphopoda | NC_005840 |
| *Graptacme eborea* | Mollusca/Scaphopoda | NC_006162 |
| *Venerupis philippinarum* | Mollusca/Bivalvia | NC_003354 |
| *Mytilus edulis* | Mollusca/Bivalvia | NC_006161 |
| *Lampsilis ornata* | Mollusca/Bivalvia | NC_005335 |
| *Inversidens japonensis* | Mollusca/Bivalvia | AB055624 |
| *Loxocorone allax* | Entoprocta | NC_010431 |
| *Flustrellidra hispida* | Bryozoa/Ectoprocta | NC_008192 |
| *Watersipora subtorquata* | Bryozoa/Ectoprocta | NC_011820 |
| *Bugula neritina* | Bryozoa/Ectoprocta | NC_010197 |
| *Paraspadella gotoi* | Chaetognatha | NC_006083 |
| *Spadella cephaloptera* | Chaetognatha | NC_006386 |
| *Sagitta enflata* | Chaetognatha | NC_013814 |
| *Sagitta naga* | Chaetognatha | NC_013810 |
| *Leptorhynchoides thecatus* | Rotifera/Acanthocephala | NC_006892 |
| *Caenorhabditis elegans* | Nematoda | NC_001328 |
| *Trichinella spiralis* | Nematoda | NC_002681 |
| *Priapulid caudatus* | Priapulida | NC_008557 |
| *Epiperipatus biolleyi* | Onychophora | NC_009082 |
| *Limulus polyphemus* | Arthropoda/Xiphosura | NC_003057 |
| *Stegancarus magnus* | Arthropoda/Arachnida | NC_011574 |
| *Dermatophajoides pteronyssinus* | Arthropoda/Arachnida | NC_012218 |
| *Leptotrombidium akamushi* | Arthropoda/Arachnida | NC_007601 |
| *Nymphon gracile* | Arthropoda/Pycnogonida | NC_008572 |
| *Narceus annularis* | Arthropoda/Myriapoda | NC_003343 |
| *Ligia oceanica* | Arthropoda/Crustacea | NC_008412 |
| *Argulus americanus* | Arthropoda/Crustacea | NC_005935 |
| *Speleonectes tulumensis* | Arthropoda/Crustacea | NC_005938 |
| *Tigriopus japonicus* | Arthropoda/Crustacea | NC_003979 |
| *Vargula hilgendorfii* | Arthropoda/Crustacea | NC_005306 |
| *Eriocheir sinensis* | Arthropoda/Crustacea | NC_006992 |
| *Megabalanus volcano* | Arthropoda/Crustacea | NC_006293 |
| *Cherax destructor* | Arthropoda/Crustacea | NC_011243 |
| *Pagurus longicarpus* | Arthropoda/Crustacea | NC_003058 |
| *Chinkia crosnieri* | Arthropoda/Crustacea | NC_011013 |
| *Balanoglossus carnosus* | Enteropneusta | NC_001887 |
| *Xenoturbella bocki* | Xenoturbellida | NC_008556 |
| *Florumetra serratissima* | Echinodermata/Crinoidea | NC_001878 |
| *Antedon mediterranea* | Echinodermata/Crinoidea | NC_010692 |
| *Gymnocrinus richeri* | Echinodermata/Crinoidea | NC_007689 |
| *Asterina pectinifera* | Echinodermata/Asteroidea | NC_001627 |
| *Ophiura lukteni* | Echinodermata/Ophiuroidea | NC_005930 |
| *Ophiopholis aculeata* | Echinodermata/Ophiuroidea | NC_005334 |
| *Strongylocentrotus purpuratus* | Echinodermata/Echinoidea | NC_001453 |
| *Doliolum nationalis* | Chordata/Tunicata | NC_006627 |
| *Phallusia fumigata* | Chordata/Tunicata | NC_009834 |
| *Phallusia mammillata* | Chordata/Tunicata | NC_009833 |
| *Ciona savignyi* | Chordata/Tunicata | NC_004570 |
| *Ciona intestinalis* | Chordata/Tunicata | NC_004447 |
| *Halocynthia roretzi* | Chordata/Tunicata | NC_002177 |
| *Assymetron inferum* | Chordata/Cephalochordata | NC_009774 |
| *Homo sapiens* | Chordata/Craniata | NC_012920 |

A comparison of the distances obtained by the approaches based on breakpoint or inversion and those obtained from this study showed the inaccuracy of these previous methods (Table 2). At a first glance, one should expect the inversion distance and the breakpoint number to be correlated with the exact minimal distance. Comparisons of lines 1 *vs* 2 and lines 3 *vs* 4 of the Table 2 shows that the number of breakpoints between two genomes can be identical while their exact minimal distance is very different. Other cases are even more surprising: the relation between exact minimal distance and the number of breakpoints moves in the opposite direction: in several examples of mtDNA pairwise comparisons (lines 5 *vs* 6, 7 *vs* 8, 9 *vs* 10 and 11 *vs* 12), the greatest number of breakpoints coincides with the shortest exact minimal distance. In several cases, the greatest inversion distances still coincide with the shortest exact minimal distances (lines 3 *vs* 4, 5 *vs* 6 and 7 *vs* 8). Moreover, while inversion distance gave a good estimation of the exact minimal distance in a few cases, it usually strongly overestimated the latter. For instance, in line 5 of Table 2, both distances are adequate because the exact minimal distance can be explained by 3 inversion events (1 path among 224 possible paths). Overestimation is generally significant when the possible minimal paths were mostly represented by transpositions. For instance, in line 3 of Table 2, the inversion distance is 9 while the exact minimal distance is 3.

**Table 2.** Comparison of some of the breakpoint number, inversion distance, and exact minimal distance results obtained from this study.

|  | Pairwise comparison of mtDNAs | Breakpoint number | Inversion distance | Exact minimal distance |
| --- | --- | --- | --- | --- |
| # 1 | *Argulus americanus* vs *Urechis caupo* | 11 | 10 | 6 |
| # 2 | *Nymphon gracile* vs *Strongylocentrotus purpuratus* | 11 | 8 | 4 |
| # 3 | *Balanoglossus carnosus* vs *Strongylocentrotus purpuratus* | 9 | 9 | 3 |
| # 4 | *Nymphon gracile* vs *Loxocorone allax* | 9 | 7 | 5 |
| # 5 | *Eriocheir sinensis* vs *Katharina tunicata* | 5 | 3 | 3 |
| # 6 | *Homo sapiens* vs *Strongylocentrotus purpuratus* | 6 | 6 | 2 |
| # 7 | *Nymphon gracile* vs *Trichinella spiralis* | 8 | 7 | 4 |
| # 8 | *Balanoglossus carnosus* vs *Asterina pectinifera* | 9 | 8 | 3 |
| # 9 | *Speleonectes tulumensis* vs *Leptotrombidium akamushi* | 9 | 8 | 5 |
| # 10 | *Limulus polyphemus* vs *Asterina pectinifera* | 10 | 7 | 4 |
| # 11 | *Ligia oceanica* vs *Graptacme eborea* | 15 | 12 | 6 |
| # 12 | *Ligia oceanica* vs *Siphonodentalium lobatum* | 14 | 13 | 7 |

Interestingly, the minimal distances calculated between mtDNAs with a maximum of 15 genes were never greater than 8, suggesting a saturation phenomenon. To highlight a putative saturation threshold, we carried out empirical tests using the simulation described below. Three datasets of 20 different genome organisations with 15, 14 and 13 genes respectively were randomly generated. In the dataset containing mtDNA with 15 genes, the minimal distances obtained for every pairwise comparison were 6 (in 15% of all the pairwise comparisons) or 7 (85%). When doing a new simulation with random mtDNAs containing 14 genes the minimal distances obtained were 6 (50%) or 7 (50%). With random mtDNAs containing 13 genes the minimal distances obtained were 5 (5%) or 6 (95%). This shows that in the case of bilaterians (with mtDNAs containing 13, 14 or 15 protein-coding and rRNA genes) if the minimal distance is greater than 5, the probability of underestimating the true number of mutations between two genomes is high. The characterisation of the saturation threshold allowed us to define a coefficient of saturation *C* for each taxon *X* with the following formula:

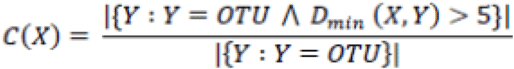

The coefficient of saturation provided an objective criterion to remove fast-evolving taxa with a coefficient of saturation close to 100% (cut off 95%) from the taxonomic dataset. Hence, 21 of 69 taxa were excluded from the analysis: all urochordates (*Ciona intestinalis*, *Ciona savignyi*, *Doliolum nationalis*, *Halocynthia roretzi*, *Phallusia fumigata*, *Phallusia mammillata*), one copepod (*Tigriopus japonicus*), one nematode (*Caenorhabditis elegans*), all scaphopods (*Graptacme eborea*, *Siphonodentalium lobatum*), all lamellibranchs (*Inversidens japanensis*, *Lampsilis ornata*, *Mytilus edulis*, *Venerupis philippinarum*), all platyhelminths (*Gyrodactylus derjavinoides*, *Schistosoma mansoni*), one acanthocephalan (*Leptorhynchoides thecatus*), two bryozoans (*Flustrellidra hispida*, *Watersipora subtorquata*) and two brachiopods (*Laqueus rubellus*, *Lingula anatina*).

### Reconstruction of mtDNA mutational networks: the deuterostomes as a study case

Different combinations of outgroups were assessed to explore their effects on the attachment point to the root and examine if the rooting choice would affect the optimal mtDNA mutational network(s). First, we computed mutational networks of deuterostomes rooted by *Limulus polyphemus* (Arthropoda, Ecdysozoa). Then, we performed analyses rooted with one or two additional taxa, (*Limulus polyphemus*, *Katharina tunicata* – Mollusca, Lophotrochozoa) and (*Limulus polyphemus*, *Katharina tunicata*, *Tethya actinia* – Porifera). All the computations are detailed in a logbook (S2 appendix). As the nature of the outgroup did not change the topologies of the optimal tree solutions, we decided to base the mutational networks discussed below on the solutions obtained from the computation #1 rooted on *Limulus polyphemus*.

The computation resulted in only six distinct solutions (S3 appendix, section ‘deuterostomes_taxA_v2_6sol’) in which the internal deuterostomes relationships are identical except some variations within Echinodermata. These variations include three topologies for the Crinoidea combined with two topologies for the rest of the Echinodermata (Fig 1). The two topologies obtained for the rest of Echinodermata consisted of different positioning of *Asterina pectinifera* and *Strongylocentrotus purpuratus,* of which the mtDNAs corresponded, in each solution, to Ur-echinodermata. This contradicts the conclusion drawn by Perseke et al. [44] who proposed that the mtDNA ground pattern of the Ophiuroidea, Crinoidea, and the group of Echinoidea, Holothuroidea, and Asteroidea could be derived from a hypothetical ancestral crinoid gene order.

**Fig 1.**
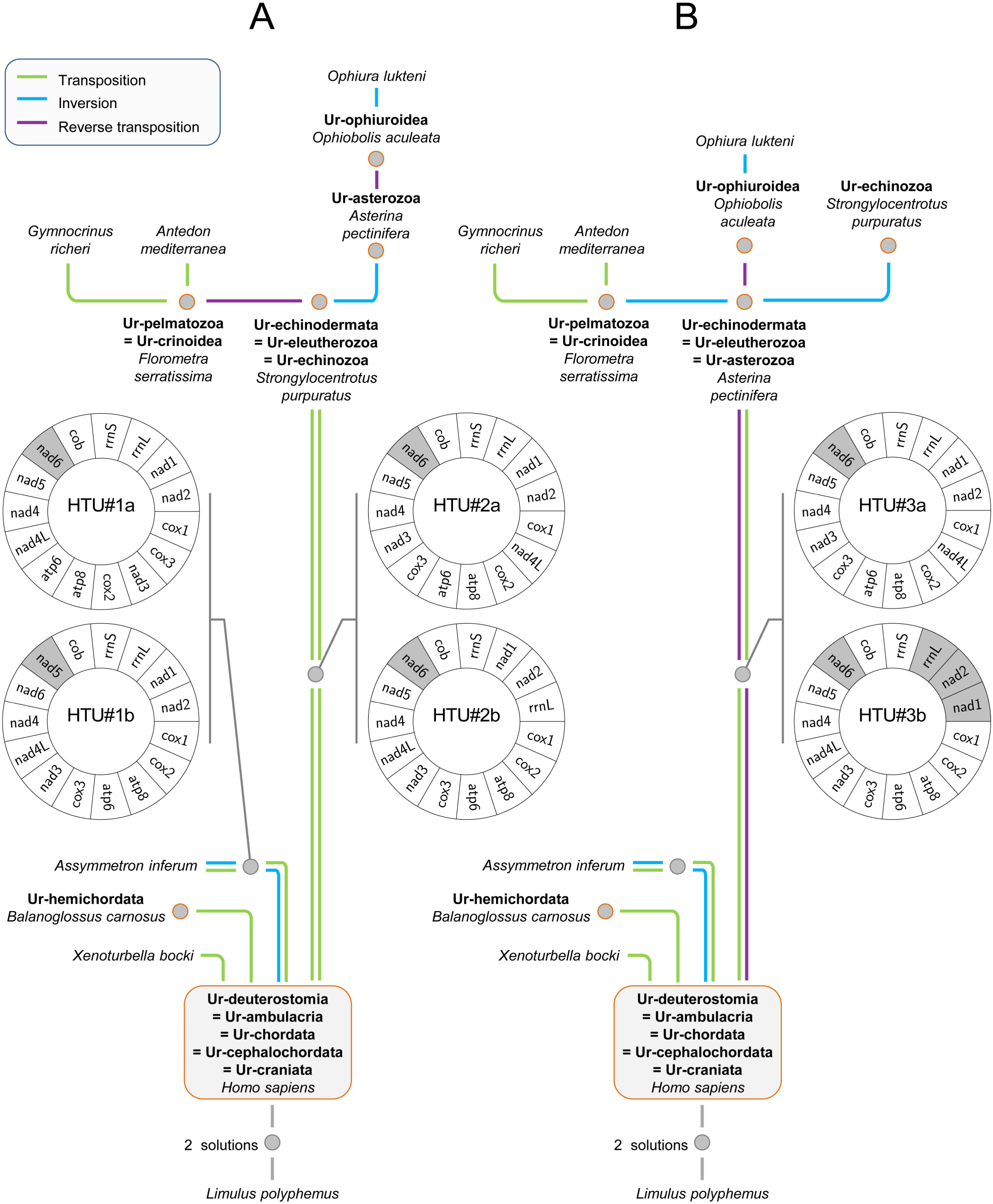
The two most parsimonious mutational networks (12 steps) based on the arrangement of protein-coding and ribosomal RNA genes in the deuterostomes mtDNAs. The maximum parsimony mutational links among the networks are indicated by solid lines (blue, inversion; green, transposition; purple, reverse transposition). Hypothetical ancestral mtDNAs (HTUs) are indicated at each node of the trees by grey dots. Orange-circled dots correspond to ground patterns of clades. Ur-echinodermata is represented by the mtDNA of either *Strongylocentrotus purpuratus* (A) or *Asterina pectinifera* (B). Grey-shaded boxes on diagrammatic representations of hypothetical ancestral mtDNAs (HTU#1a to 3b) highlight genes transcribed from the opposite strand.

In an attempt to further explore the mutational network of extant echinoderms, reduce the number of solutions, and specify the mtDNA organisation of Ur-echinodermata, we used three additional and alternative PPHs defined from the most recent literature on Echinodermata:

- PPH#30a (Echinoidea+Asteroidea) [44]. In this first hypothesis, the Asteroidea are placed as a sister group to the clade Echinoidea + Holothuroidea
- PPH#30b (Ophiuroidea + Echinoidea): Cryptosyringid [45]. In this second hypothesis, the Ophiuroidea are placed as a sister group to the clade Echinoidea + Holothuroidea
- PPH#30c (Asteroidea + Ophiuroidea): Asterozoa [46]. In this third hypothesis, the Asteroidea are placed as a sister group to Ophiuroidea.

When tree reconstructions were constrained with PPH#30a or PPH#30b we still found three topologies corresponding to three different relationships within Crinoidea, but *Asterina pectinifera* always appeared as Ur-echinodermata (Fig 1B), whatever the PPH considered. Only with PPH#30c, computations show that either *Asterina pectinifera* or *Strongylocentrotus purpuratus* represents Ur-echinodermata in distinct but equiparsimonious scenarios (Fig 1). Previous analyses based on mt gene order have suggested that the ancestral mt genome of echinoderms most likely resembles the echinoid mtDNA [13, 47]. However, a logical approach showed that either Echinoidea or Asteroidea might represent the echinoderm ground pattern.

Three topologies exhibited different relationships within Crinoidea, specifically between *Antedon mediterrannea* and *Gymnocrinus richeri* with *Florometra serratissima* basal to Crinoidea (S3 appendix, section ‘deuterostomes_taxA_v2_6sol’). For a local analysis of the mutational network within Crinoidea, tRNA genes have been included for computations. There are up to 22 tRNA genes added to the 15 protein-coding and rRNA genes. Therefore, including the tRNAs into the path calculation between two genomes increases the computation time to a point that makes the procedure unfeasible, unless the genomes compared are very close. For example, the path calculation between *Florometra serratissima* and *Antedon mediterrannea* is fast, as the minimal distance is 3. However, between *Asterina pectinifera* and *Homo sapiens,* the minimal distance is at least 11, whereas it is only 2 when considering only protein-coding genes. When *Asterina pectinifera* or *Strongylocentrotus purpuratus* are used as outgroups, the analysis that included the tRNA genes yielded a single unique topology for the Crinoidea (S3 appendix, sections ‘with_tRNA_crinoids_taxA_1sol’ and ‘with_tRNA_crinoids_taxB_1sol’), as represented in Fig 1.

Finally we performed computations regarding several phylogenetic hypotheses on *Xenoturbella bocki* relationships. Acoelomorph flatworms related to *Xenoturbella bocki* were initially placed within deuterostomes [48] but several conflicting hypotheses are still under debate. A first study based on the analysis of newly sequenced mtDNAs [49] provided no support for a sister group relationship between Xenoturbellida and Acoela or Acoelomorpha and suggested an unstable phylogenetic position of *Xenoturbella bocki* as sister group to or part of the deuterostomes. More recently, two phylogenomic analyses have grouped *Xenoturbella* with acoelomorphs (=Xenacoelomorpha) and suggested that Xenacoelomorpha could be the sister group of Nephrozoa [50, 51] or Protostomia [50]. In our contribution, whatever the PPH used to constrain the position of *Xenoturbella bocki* (*i.e*., whether it is sister group to or part of the deuterostomes), its mtDNA organisation is always derived from an ancestor exhibiting an mtDNA organisation identical to *Homo sapiens* (S3 appendix). Hence our results corroborate the conclusion that the arrangement of protein-coding and rRNA genes in the mtDNA of *Xenoturbella bocki* is plesiomorphic [52] and therefore does not contain relevant signal to assess the phylogenetic relationships of this species.

As a result of using PPHs to constrain the relationships within echinoderms and tRNA genes to decipher Crinoidea relationships, only two mutational networks were finally validated for deuterostomes (Fig 1). The major advantage of a complete approach is that all the values of HTUs (inferred ancestral mtDNA organisations) that are by definition not present in the taxonomic dataset are enumerated. Such a comprehensive and correct enumeration is not possible in traditional probabilistic approaches or by manual inspection of pairwise scenarios. In the case of the deuterostomes, calculating the mtDNA mutational network required the insertion of only two HTUs, the first in the lineage leading to cephalochordates and the second in the one leading to the echinoderms (Fig 1). Each HTU has two possible values because of the commutative property of both paths described. For each path, the HTUs represent a ground pattern that characterizes an ancestor or a current mtDNA that has not been sequenced yet. Interestingly, the mt gene orders of HTU#2a and HTU#3a that stand for two distinct paths between Craniata and Echinodermata have already been characterised in a previous study and were considered as the echinoderm consensus [47].

To summarise, below are listed the key results from this first analysis, results that could also be described as logical consequences of the problem PHYLO:

- Monophyly of Chordata, monophyly of Echinodermata, monophyly of Ophiurida and monophyly of Crinoidea are always verified.
- There is only one subtree for Cephalochordata with *Homo sapiens* mtDNA as Ur-cephalochordate.
- There is only one subtree for Ophiuroidea with *Ophiobolis aculeata* mtDNA as Ur-ophiuroidea.
- There is only one position for Hemichordata and *Xenoturbella bocki.*
- There is only one subtree for Crinoidea (when using tRNA genes) and *Florometra serratissima* mtDNA represents Ur-crinoidea.
- *Homo sapiens* mtDNA represents Ur-deuterostomia, Ur-chordata, Ur-cephalochordata and Ur-ambulacria.

### The use of tRNA genes to solve the PHYLO problem

tRNA genes are usually omitted in the comparison of mt gene arrangements because of their accelerated moving rate. However, as shown above, tRNA genes do contain phylogenetic information in some contexts and should be considered in rearrangement models to limit the number of tree solutions. The computation of the Echinodermata mutational network with all mt genes was possible but could not be held with completeness. The reconstruction of one tree solution with all mt genes with *Asterina pectinifera* as Ur-echinodermata (S3 appendix, sections ‘with_tRNA_eleutherozoa_taxA_2sol’ and ‘with_tRNA_ophiurida_taxA_1sol’) required 25 evolutionary steps and was tractable (Fig 2A). This topology was slightly different (different branching within Ophiuroidea) from the topology obtained with the protein-coding and rRNA genes on which specific mutations of tRNA genes have been added *a posteriori* (Fig 2B). The ancestral state of Ophiuroidea has been shown to be difficult to infer and remains unresolved [53, 54] but it has been suggested that *Ophiura lutkeni* has a more derived mt gene order than *Ophiobolis aculeata* [53]. While the scenarios computed with protein-coding and rRNA genes always favoured *Ophiobolis aculeata* mt gene order as the Ophiuroidea ground pattern (Fig 1), the topology obtained with the inclusion of tRNA genes proposed 6 distinct hypothetical ground patterns (Fig 2A) with a more derived position for *Ophiobolis aculeata* than *Ophiura lutkeni*. Moreover, a TDRL event has been proposed in the path between Echinoidea and Holothuroidea [54, 55]. In the present work, all the 14,641 possible path of length five between *S. purpuratus* and *C. miniata* are represented by a succession of five tRNA transpositions (Fig 2).

**Fig 2.**
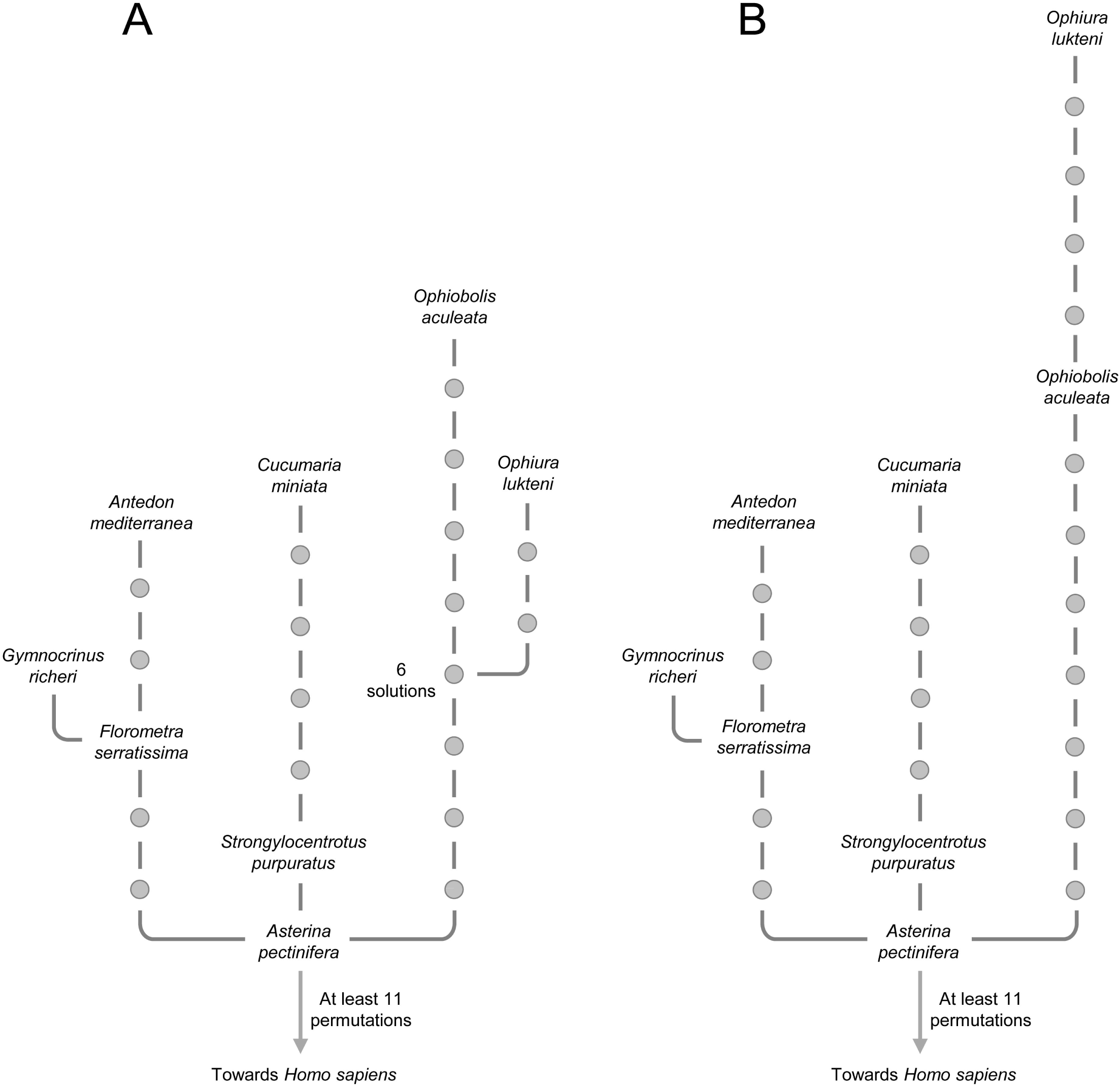
Mutational networks based on the arrangement of all the mt genes in echinoderms. (A) One tree solution for the whole Echinodermata group showing a possible mutational network calculated with all mitochondrial genes (including tRNA genes). Among the 25 necessary steps, more than 6 involve the mitochondrial coding-protein and rRNA genes. (B) Tree solution calculated with mitochondrial protein-coding and rRNA genes with *Asterina pectinifera* as Ur-echinodermata (Fig 1B) on which 20 necessary permutations of tRNA genes have been added *a posteriori*. Among the 26 steps, 6 involve the mitochondrial coding-protein and rRNA genes.

The computation including all mt genes (Fig 2A) raised interesting remarks. In the mutational network computed with all mt genes, the total number of permutations was minimal and represented the most parsimonious scenario (25 permutations on Fig 2A instead of 26 on Fig 2B obtained from a computation with coding-protein and rRNA genes only). However the total number of evolutionary steps concerning the protein-coding genes increased (Fig 2A, always more than 6 permutations) when compared with the least parsimonious scenario computed without tRNA genes (Fig 2B, 6 permutations). Parsimony is the principle according to which, all other things being equal, the best hypothesis to consider is the one that requires the fewest evolutionary changes. However, the reasonableness of the parsimony rule in a given context may have nothing to do with its reasonableness in another one. In other words, when using the parsimony principle to decipher evolutionary hypotheses, the outcome depends on the set of characters considered. Indeed, when we look at the history of a given mtDNA, nearly 80% of all the permutations that have happened involve tRNA genes. Given this high percentage, if we want to minimize the global number of permutations (*i.e*., if we are looking for parsimonious trees that takes all gene into account), the influence of larger protein-coding and rRNA genes is negligible when compared to the one of smaller tRNA genes. Hence, the mutational networks obtained only with the larger genes are expected to be significantly different than those obtained with all the genes (which should be very similar to the parsimonious trees obtained when using only tRNA genes). This suggests that even if the use of tRNA genes can be relevant in certain cases for local resolutions, it is reasonable to rely predominantly on the larger genes with a lower evolutionary rate, when calculating the tree solutions corresponding to deep and ancient nodes/lineages like in the case of deuterostomes or bilaterians.

### Towards the mtDNA mutational network of bilaterians with a logical approach

There were too many OTUs (47 bilaterians + 1 poriferan) to make a single global computation, but smaller computations that verify the convergence of results at each step were tractable.

Using known monophyletic groups (see Methods and S9 appendix), calculations were carried out on taxonomic groups and subgroups by recombining the resulting solutions in the hierarchical structure of the Bilateria phylogeny. The chronological description of all the computations is given in S2 appendix. A high number of equiparsimonious topologies were obtained. Even though a unique representation of these topologies is not possible, the whole set of solutions can be enumerated (S3-S6 appendix). The solution files make up a database of all the possible solutions. Moreover, as described in the example above with the echinoderms, the number of possible solutions can be reduced, possibly down to a single one, by adding PPHs to the calculation.

In the case of Ecdysozoa, seven calculations had to be carried out to obtain a complete mutational network (S4 appendix). After the recombination of these calculations, 4212 equiparsimonious solution trees were obtained corresponding to 3 subtrees for Decapoda combining with 39 subtrees for the rest of Mandibulata, (3×39 = 117 subtrees for all Mandibulata), 9 subtrees for Chelicerata (comprising 6 subtrees for Acari, meaning 117×9 = 1053 subtrees for Arthropoda), 1 subtree for Onychophora (1×1053 = 1053 subtrees for Panarthropoda), 4 subtrees for Introverta (4×1053= 4212 trees for Ecdysozoa).

Concerning Lophotrochozoa, 12 calculations were needed, leading to 81 solution trees (S5 appendix): 3 subtrees for Gasteropoda combining with 1 subtree for the rest of Mollusca (1×3= 3 subtrees for Mollusca), 3 subtrees for the rest of Eutrochozoa, (3×3 = 9 for Eutrochozoa), 9 subtrees for Lophophorata (9×9 = 81 subtrees for Lophotrochozoa).The large amount of equiparsimonious solution trees obtained for the two main protostomian clades does not allow a single representation. Nevertheless, the analyses provided several logical consequences that are important results from a biological perspective:

- Monophyly of Acari, monophyly of Panarthropoda, and monophyly of Annelida are always verified.
- *Limulus polyphemus* mtDNA represents Ur-panarthropoda, Ur-arthropoda, Ur-mandibulata and Ur-chelicerata.
- There is only one solution for the set of mutations which links the mtDNA of Ur-panarthropoda (*Limulus polyphemus*) and the mtDNA of *Eriocheir sinensis* (transposition), *Narceus annularis* (transposition) and *Epiperipatus biolleyi* (6 possible paths, each with 3 mutations).
- Three solutions have been found for Ur-ecdysozoa which correspond either to the mtDNA of *Limulus polyphemus* or to *Priapulus caudatus* or to a hypothetical ancestor (HTU#1).
- There is only one solution for the cephalopod clade, with *Katharina tunicata* mtDNA as Ur-cephalopods linking *Nautilus macrocephalus* mtDNA (transposition) and *Loligo bleekeri* mtDNA (4 possible paths, each with 2 mutations).
- *Cepaea nemoralis* represents Ur-gasteropoda.
- There is only one solution for the position of *Loxocorone allax* mtDNA (10 possible paths, each with 2 mutations) and *Phoronis architecta* mtDNA (one transposition) with respect to *Katharina tunicata.*
- *Sipunculus nudus* (Sipunculida) is always the sister group of all the annelid species (*Platynereis dumerilii* and *Urechis caupo*).
- *Katharina tunicata* mtDNA represents Ur-lophotrochozoa, Ur-eutrochozoa, Ur-mollusca, Ur-lophophorata and Ur-cephalopoda.

To give more insight into the deep branching of bilaterians, we carried out a computation rooted on *Tethya actinia* and using the respective gene order ground patterns of protein-coding and rRNA genes of ecdysozoans (*Limulus polyphemus*, *Priapulus caudatus* or HTU#1), lophotrochozoans (*Katharina tunicata*) and deuterostomes (*Homo sapiens*) as the representative of the three main bilaterian clades. This strategy allowed enumerating with completeness 6 mutational networks for Bilateria (Fig 3, Table 3) and to highlight the following logical consequence: *Homo sapiens* mtDNA represents Ur-bilateria.

**Fig 3.**
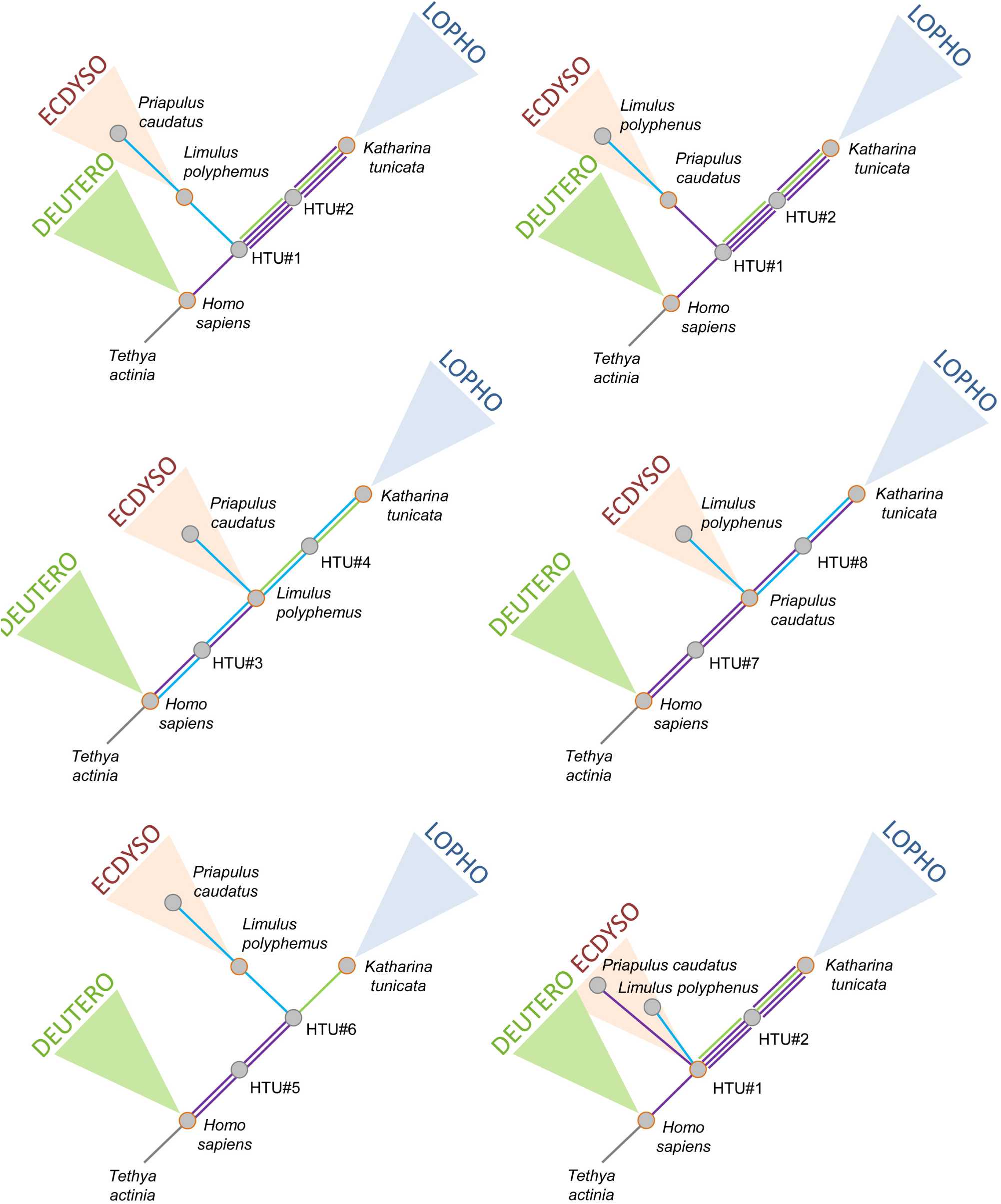
The six most parsimonious mutational networks based on the rearrangements of protein-coding and rRNA genes in bilaterian mtDNAs. The maximum parsimony mutational links among the networks are indicated by solid lines (blue, inversion; green, transposition; purple, reverse transposition). Hypothetical ancestral mtDNAs are indicated at each node of the trees by grey dots. Orange-circled dots correspond to ground patterns of deuterostomes, ecdysozoans and lophotrochozoans. Orders of mitochondrial protein-coding and rRNA genes of HTU#1 to 8 are given in Table 3.

**Table 3.** Organisation of mitochondrial protein-coding and ribosomal RNA genes of hypothetical ancestral mitochondrial genomes (HTUs) represented on Fig 3.

| HTU # | Hypothetical genomic organizations |
| --- | --- |
| 1, 3, 5, 7 | cox1 cox2 atp8 atp6 cox3 nad3 -nad5 -nad4 -nad4L nad6 cob rrnS rrnL nad1 nad2 |
| 2 | cox1 cox2 atp8 atp6 -nad5 -nad4 -nad4L nad6 cob rrnS rrnL nad1 cox3 nad3 nad2 |
| 2 | cox1 cox2 atp8 atp6 -nad5 -nad4 -nad4L nad6 cob -nad3 -cox3 rrnS rrnL nad1 nad2 |
| 2 | cox1 cox2 atp8 atp6 nad6 cob nad4L nad4 nad5 -nad3 -cox3 rrnS rrnL nad1 nad2 |
| 2, 4, 6 | cox1 cox2 atp8 atp6 cox3 nad3 -nad5 -nad4 -nad4L -cob -nad6 -nad1 -rrnL -rrnS nad2 |
| 3 | cox1 cox2 atp8 atp6 cox3 nad3 nad4L nad4 nad5 -nad6 cob -nad1 -rrnL -rrnS nad2 |
| 4, 8 | cox1 cox2 atp8 atp6 -nad5 -nad4 -nad4L nad6 cob -nad1 -rrnL -rrnS cox3 nad3 nad2 |
| 5 | cox1 cox2 atp8 atp6 cox3 nad3 -nad5 -nad4 -nad4L -cob nad6 rrnS rrnL nad1 nad2 |
| 7 | cox1 cox2 atp8 atp6 cox3 nad3 rrnS rrnL nad1 -cob nad6 -nad5 -nad4 -nad4L nad2 |
| 8 | cox1 cox2 atp8 atp6 cox3 nad3 rrnS rrnL nad1 nad6 cob nad4L nad4 nad5 nad2 |

Mt gene arrangements and ground patterns in Bilateria have been previously studied in Lophotrochozoa [28, 56], Ecdysozoa [57] and Deuterostomia [52]. It is important to note that these studies usually considered all the mt genes to draw their conclusions, which could explain some incongruence with the present results. Notably, it has been suggested that the ancestral mt gene order in Lophotrochozoa and Deuterosotomia cannot be found in extant species but rather represent consensus between ingroup and outgroup arrangements [28]. Considering the protein-coding and rRNA genes, we showed that the ground patterns of Deuterostomia and Lophotrochozoa are realized in the respective mt gene arrangements of two extent species, *Homo sapiens* and *Katharina tunicata.* In Ecdysozoa, Ur-arthropoda is always realized in the gene arrangement of *Limulus polyphemus* like it has been previously proposed [58]. In addition, the mt gene order of *Limulus polyphemus* should also be considered as the ground pattern of Panarthropoda and Ecdysozoa but our results also demonstrated that Ur-ecdysozoa could also correspond to the mitochondrial genome of *Priapulus caudatus* or in an inferred ancestral mtDNA organisation that is not realized in extant species. Priapulids have been described as an ancient clade and seem likely to adhere closely to the predicted ecdysozoan ground pattern [57]. Finally, the ground pattern of Bilateria was previously hypothesised [59]. The arrangement of the protein-coding and rRNA mt genes of *Homo sapiens* has been considered as Ur-bilateria. It has been also suggested that the differences previously observed between vertebrate and arthropod genomes are due mainly to gene rearrangements within the protostome lineages, a conclusion corroborated by our study.

Additional computations rooted on *Tethya actinia* were carried out with four chaetognath mtDNAs added to the dataset described above (S7-S8 appendix). The position of chaetognaths was either basal to protostomes, ecdysozoans, or lophotrochozoans (34 possible topologies, S7-S8 appendix) and three logical consequences were emphasised:

- Chaetognatha mtDNAs were always grouped together (monophyly of Chaetognatha is always verified).
- Among the chaetognaths, the Sagittidae family is valid with *Flaccisagitta enflata* mtDNA as Ur-sagittidae.
- Chaetognatha mtDNAs cannot be basal to all bilaterians (the mtDNA organisation of chaetognaths never derived directly from that of *Homo sapiens*).

Although it was possible to assert that chaetognaths were not the sister group of bilaterians, the different topologies obtained are another reminder that the phylogenetic position of Chaetognatha is still one of the most problematic issues of metazoan phylogeny [60].

## Conclusions

We hope that this pilot work will inspire further use of formal logic for evolutionary studies based on gene order data. It has the benefit of both correctness and completeness: in our study, the tree solutions obtained for each computation encompass all the mutational networks that verify properties P1 to P6. It is impossible to explore all the entire possible solutions by manual inspection when the path length is more than one step. Such an approach is non-exhaustive and therefore useless. At first, the exploration of all the possible mutational networks might not seem to be a very elegant method, as it can offer numerous solutions to the same problem. However, an understanding of the logical consequences (true for all solutions found) can only be obtained through a complete enumeration and these logical consequences are, in themselves, extremely robust results. In our study of the bilaterian mtDNAs, we have decided to use the broadest and most indisputable PPHs, which lead us to all the possible mutational networks and, indeed, to a high number of equiparsimonious trees. By adding more PPHs for higher-level bilaterian taxa, we have significantly reduced the number of solutions depending on the supplementary hypotheses selected. Such a hypothetico-deductive approach was particularly fruitful in the study case of deuterostomes and should be applied to many other clades of bilaterians.

## Methods

### Problematic

The dataset is represented by a taxonomic sample of *N* metazoans for which the mtDNA is known and a set of possible elementary mutations to switch between one mtDNA to another. It is possible to encode the organisation of a given mtDNA by a linear representation that satisfies the following properties:

- the last gene in the list precedes the first one (mtDNA circularity);
- when the sign of cox1 is positive (positive strand), the gene list is read from left to right;
- when the sign of cox1 is negative (negative strand) the gene list is read in the reverse direction and the signs of all the other genes are inverted.

From a linear representation and, if necessary, by reversing the reading direction and the sign of genes it is possible to define a Canonical Linear Representation (CLR) for all mtDNAs with the following properties:

- the first gene is cox1 (by convention);
- the sign of cox1 is positive and genes are read from left to right.

For instance the CLR of *Homo sapiens* mtDNA with thirteen protein-coding and two ribosomal RNA (rRNA) genes is given by:

[cox1 cox2 atp8 atp6 cox3 nad3 nad4L nad4 nad5-nad6 cob rrnS rrnL nad1 nad2]

#### Definition 1: successive genes

Let *A* be a genome and *g*_1_ and *g*_2_ be two genes in *A*. We can say that *g*_1_ and *g*_2_ are successive in *A* if *g*_2_ follows *g*_1_ in the CLR of *A* or if *g*_1_ is the last gene in the CLR of *A* and *g*_2_ the first gene in the CLR of *A* (*g*_2_ being cox1).

#### Definition 2: block of genes

Let *A* be a genome. A block of genes is a sequence of successive genes [*g*_1_ *g*_2_…*g_r_*] in *A* (*g_i_* and *g*_*i*+*1*_ are successive in *A*).

The set of five possible elementary permutations consists of:

- transposition: a gene or a block of genes (named “selected block”) separates from the genome and is re-inserted between two genes at a different position (named “insertion point”).
- inversion: a gene or a block of genes separates from the genome and is re-inserted in the opposite direction at the same position. When a block of genes reverses, the gene order within the block is reversed as well as the sign of each gene.
- reverse transposition: a transposition in which the re-inserted gene or block of genes is reversed;
- loss: a gene or a block of genes separates from the genome and disappears;
- gain: a novel gene or a block of genes is inserted in the genome.

Solving the PHYLO problem with completeness consists of enumerating all the equiparsimonious trees that explain the paths between distinct mtDNAs with the minimal number of genomic events. In order to calculate these trees, the reconstruction method is organised along three main procedures. First, a pairwise genome comparison program calculates the minimal paths between all mtDNAs encoded in a minimal distance matrix. Second, a complete finite model generator for first-order logic calculates all the most parsimonious trees that respect the minimal distance matrix and clades defined by Primary Phylogenetic Hypotheses (hereinafter PPHs). Third, ancestral mt gene arrangements (or Hypothetical Taxonomic Units, hereinafter HTUs) are defined at all internal nodes. For this study, 29 PPHs were used (S9 appendix). They all are well-admitted hypotheses associated to well-known taxa. Our results showed that 8 among these PPHs were logical consequences, *i.e*., they were always verified even not previously imposed (see Results and Discussion). Consequentially, only the 21 PPHs imposing the monophyly of the following taxa were necessary for the study: Bilateria, Deuterostomia, Ambulacria, Eleutherozoa,Ecdysozoa, Arthropoda, Mandibulata, Crustacea, Decapoda, Chelicerata, Introverta, Lophotrochozoa, Mollusca, Polyplacophora, Cephalopoda, Gasteropoda, Eutrochozoa, Polychaeta, Echiura, Lophophorata, and Brachiopoda.

#### Step 1 - Genome comparison algorithm

Let *A* and *B* be two mtDNAs. The calculation of all possible paths in *k* steps to move from *A* to *B* by successive elementary mutations is an NP-hard problem. We wrote a program genome_comparison.c (source code available upon request) that, through a depth first exploration of a search tree, enumerates all the possible paths of length *k* starting from *A*. Each path of length *k* leading to *B* is a solution while any other path (not ending to *B*) is a deadlock which causes backtracking in the search tree and exploration of another branch (this is the backtracking algorithm, see [61]). The backtracking algorithm to find a path from *A* to *B* in *k* steps is given below.

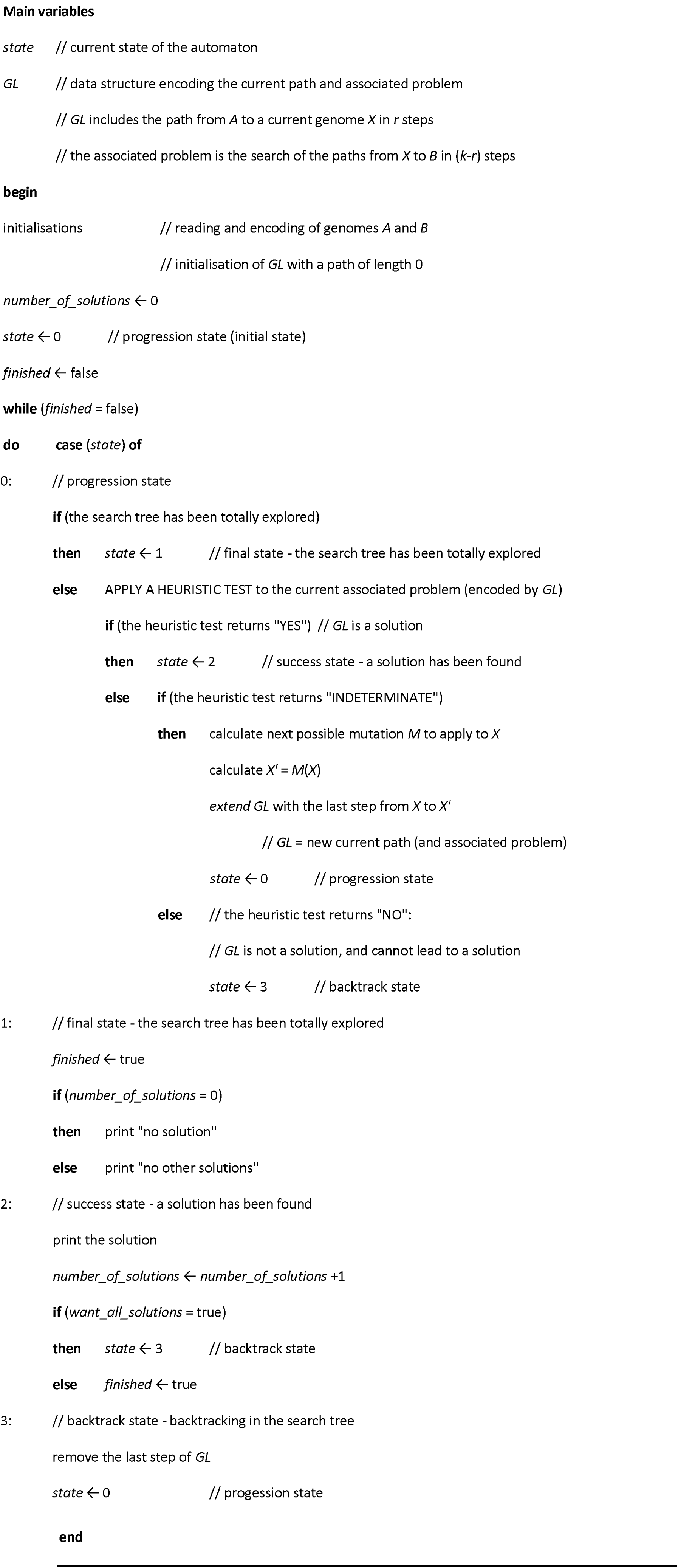
Backtracking algorithm: calculation of all the paths between *A* and *B* mtDNAs in *k* steps.

In state 0 (progression state), we apply a heuristic test as a function that returns:

- YES, if the path encoded in *GL* is a solution, *i.e*., a path from *A* to *B* in *k* steps. In this case, the solution is displayed (state 2).
- NO, if the path encoded in *GL* is not a solution and cannot be extended to construct a solution. In this case, the algorithm backtracks (state 3) and does not need to explore the search subtree from the current path *GL* because it cannot lead to a solution. This reduces the computation time.
- INDETERMINATE, otherwise. In this case, the algorithm extends the current path *GL* by listing all the possible mutations it can apply to the current genome *X*.

To calculate the next mutation to be applied to the current genome *X*, all possible mutations must be enumerated: transposition, reverse transposition, inversion and gain/loss. But it is possible to improve the program by not considering the gain/loss events and calculating the paths between two mtDNAs reduced to their common genes (with the same number of genes) only using transposition, reverse transposition and inversion. Then, and still with completeness, the model generator calculates the tree solutions that fit with the minimal distances. Because in the considered dataset, gain/loss events are rare and obvious, the solutions trees are determined discarding these events which are inserted *a posteriori*.

Due to genome circularity, there are always three distinct transpositions leading to a single gene order (Fig 4). Therefore, the program arbitrarily chooses one of these three possibilities when it enumerates transpositions.

**Fig 4.**
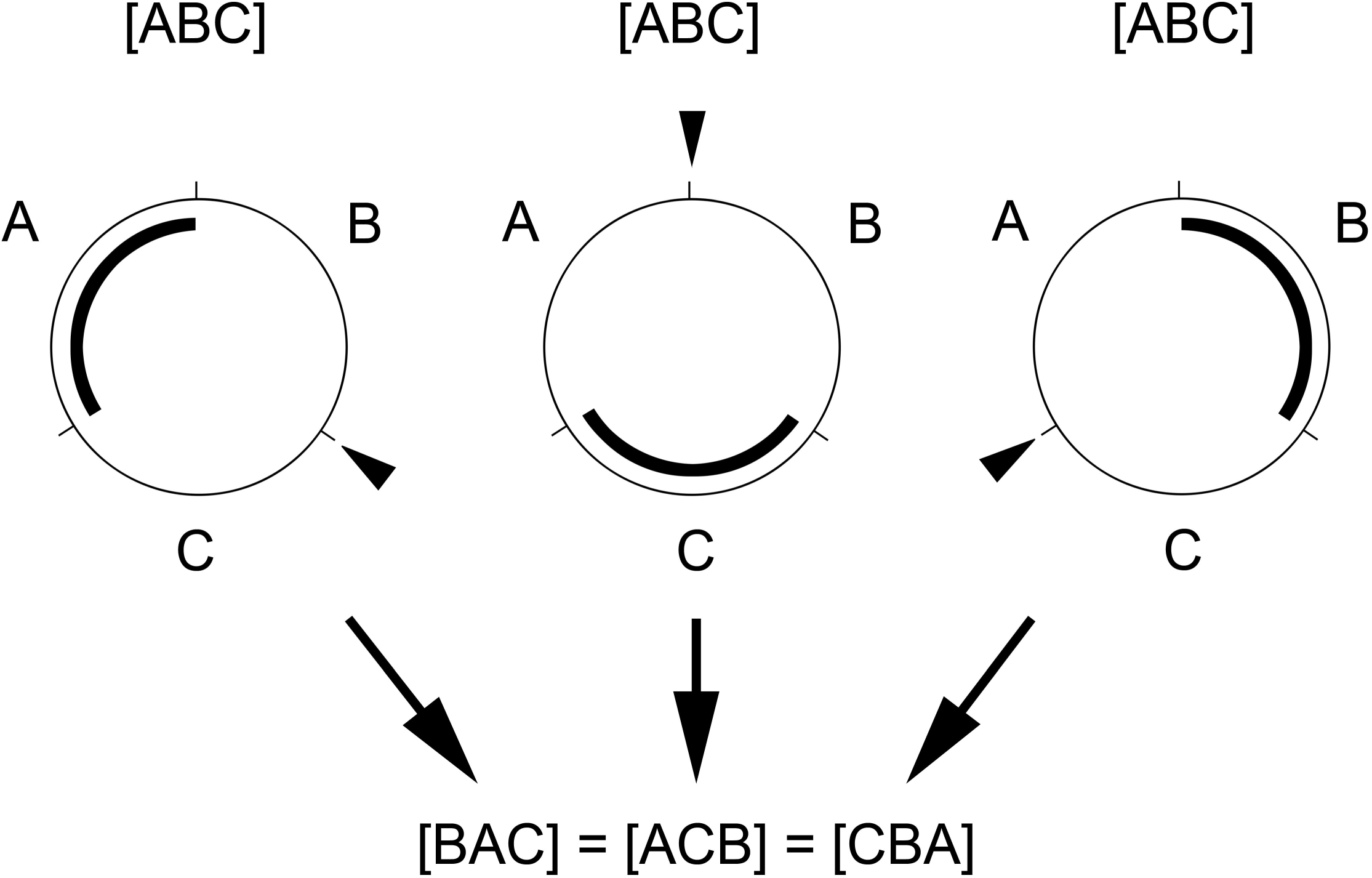
Diagram of three possible transpositions in a circular genome leading to a same gene order. Because of the circularity of the genome, there are always three possible transpositions leading to a similar gene order ([BAC] = [ACB] = [CBA]) from a given gene order ([ABC]). Thus, it is not possible to determine which gene or block of genes is concerned by a transposition. In each example, the block that is being transposed is underlined. The black arrowheads indicate where the transposed block will be inserted (insertion points).

Because of the exponential complexity of the backtracking algorithm, the associated computation time can be very long for paths with numerous rearrangement steps. This time can however be dramatically reduced when using two demonstrated mathematical properties as heuristic tests, the shared blocks property and the lower bound for minimal distances property (see Results and Discussion). Here, we present four useful definitions towards the mathematical proof of the two properties cited above:

#### Definition 3: breakpoint

Let *A* and *B* be two genomes having exactly the same genes. Let *g*_1_ and *g*_2_ be successive genes in *A*. The position (in *A*) between *g*_1_ and *g*_2_ is called a breakpoint between *A* and *B* if neither *g*_1_ and *g*_2_ nor -*g*_1_ and -*g*_2_ are successive in *B*. The number of breakpoints between *A* and *B* is denoted by *nb_breakpoints*(*A*, *B*).

#### Definition 4: shared block

Let *A* and *B* be two genomes having exactly the same genes. A block shared between *A* and *B* is a block of genes containing no breakpoint. Equivalently, a block shared between *A* and *B* is a block of genes that appears in both *A* and *B* (possibly in a reversed order)

#### Definition 5: maximum shared block

Let *A* and *B* be two genomes having exactly the same genes. A maximum shared block between *A* and *B* is a shared block that is delimited by two breakpoints.

#### Definition 6: cut

Let *C* = (*A*_0_, *A*_1_,…,*A_k_*) be a *k*-path from genome *A*_0_ to genome *A_k_*. Given an elementary mutation *μ_i_* (transposition, inversion, reverse transposition) that transforms *A_i_* into *A*_*i*+1_, *μ_i_* is called a cut if two blocks *G*_1_ and *G*_2_ exist and are successive in *A_i_* and *A_j_* (*j* > *i*+1) but not in *A*_*i*+1_. In other words, [*G*_1_ *G*_2_] is a block shared between *A_i_* and *A_j_*, but is intersected in *A_i_* by the mutation *μ_i_*.

#### Step 2 - Tree computation

Logic allows studying a problem with a hypothetico-deductive approach, permitting the enumeration of all the solutions of the problem under study, to then assess working hypotheses or answer specific questions, all with the same program (Genesereth and Nilsson, 1988) [62]. A finite model generator is a program that computes all the solutions of a set of first-order logical formulas representing the problem. Several model generators exist [63], and their common characteristics are correctness (the solutions are correct), completeness (all the solutions are listed), and decidability (all computations end, an obvious property for finite domains). PHYLO can be axiomatised by writing a set of first order logic formulas that defines a connected and acyclic graph (tree or dendrogram) representing the network of the minimum number of mutations between all the distinct mtDNAs of a given taxonomic dataset. In this graph, each vertex (node) of the tree represents an mtDNA, while each edge represents a mutation between two linked nodes.

To write this set of logical formulas (see axioms in S3-S7 appendix), one can use a relation *R*(*x*, *y*) which represents by convention the existence of an edge between the nodes *x* and *y* in the graph, and each instance of the relation *R*(*x*, *y*) being true or false. In other words, *R*(*x*, *y*) is true when there is an edge between *x* and *y*. The set of all possible values of the arguments of the relation *R* is called the domain (this is the set of all the nodes of the graph).

Finally, the solutions of PHYLO are the most parsimonious graphs *G* (*i.e*., defined on the smallest possible domain containing at least all the taxonomic dataset) which satisfy the following axioms:

**Property P1** - *G* is simple (the relation *R* is not reflexive, *i.e*., *R*(*x*, *x*) is false).

**Property P2** - *G* is non-oriented (*R* is symmetrical).

**Property P3** - *G* is connected and acyclic (*G* is a tree)

**Property P4** - *G* respects the minimal distance matrix: for each pair of genome *x* and *y* belonging to the taxonomic dataset, the path length between the nodes *x* and *y* in *G* is always greater than or equal to the minimal distance between *x* and *y*.

**Property P5** - *G* verifies additional constraints (PPHs), conditional on choosing a root node to define the hierarchical levels in the tree. In other words, the PPH imposes the existence of given monophyletic groups.

**Property P6** - It is possible to calculate all the possible mtDNA organisations for each HTU in *G* (the ancestral states that are not represented in the dataset).

The model generator computes all the solutions that satisfy the properties P1 to P5 considered as axioms of PHYLO. Property P6 is verified *a posteriori* for each solution.

In a model generator, it is possible to add extra-logical constraints to the set of formulas. These are defined an algorithmic process called constraint programming [64]. In practice, a constraint can replace a group of logical formulas having the same meaning. This can improve performance by replacing as a large subset of logical formulas maybe replaced by an equivalent constraint that is checked more easily and speedily. This feature is supported by the Davis and Putman model generator [65, 66]. At each step, the program checks if the current interpretation satisfies the constraints or not. If the constraints are not satisfied, the current interpretation is rejected and there is backtracking. For instance, the property P3 (the graph is connected and acyclic) can be easily verified for each current interpretation by a constraint that verifies that, for each connected components of the current graph, the number of nodes equals the number of edges plus 1. Similarly, the properties P4 and P5 are preferably expressed in the form of constraints rather than sets of logical formulas.

In the present work, we used an experimental model generator belonging to the Davis and Putnam type with symmetry breaking techniques [67, 68]. This model generator is correct, complete, and decidable and supports constraints. Any model generator with similar characteristics may also be suitable to solve PHYLO.

#### Step 3 - Calculating the hypothetical common ancestors (verifying P6)

After step 2, the analysis of each tree solution enables determination of ancestral states (also called ground patterns). In each tree solution with *V* nodes, there are *N* nodes (*N* < *V*) representing the *N* genomes belonging to the taxonomic dataset (Operational Taxonomic Units, OTUs), and *M* additional nodes that represent ancestral states (HTUs): *V* = *N* + *M*. Determining ancestral states consists in enumerating all possible values for the *M* HTUs of the tree solution (and thus proving that the tree solution verifies property P6), or on the contrary in proving that there is at least one HTU in the tree solution for which no value can be found (in this case the tree solution does not verify property P6, and therefore is not valid).To enumerate all the possible values of the *M* HTUs, all the paths of length *k* linking two OTUs *A* and *B* must be recalculated such that the path between *A* and *B* in the tree solution is of length *k* and passes only through HTUs (the program “genome_comparison.c” enumerates all the paths). It appears that two cases are possible for each HTU *X*:

1. A unique pair (*A*, *B*) of OTUs is such that *X* appears in the recalculated paths between *A* and *B*. In this case, all possible values for *X* appear in the paths between *A* and *B* at the position corresponding to *X.*
2. Several pairs of OTUs are such that *X* appears as an intermediate step in the recalculated paths. In this case, it is necessary to find the common values for *X*. This could be done using a simple text editor by searching for common mtDNAs in the files containing all possible paths, but to perform this operation we used a string comparison program available upon request. A lack of common values for *X* means that the tree solution is not a valid solution: it does not verify P6 because it contains a minimal subtree that does not verify P6. Any other solution containing this minimal subtree is similarly not valid.

#### Step 4 - Determining the complete set of tree solutions

For each invalid minimal subtree *A*, an additional constraint is programmed into the model generator to rule out the solutions containing *A*. Tree solutions are recalculated (step 2) and verified (step 3), possibly leading to the discovery of other invalid minimal subtrees and thus to the addition of new constraints to recalculate the solutions (feedback mechanism). The complete set of optimal solutions is determined by iterating this process and eliminating all the solutions that do not verify P6.

A result verified by all the enumerated solutions is called a logical consequence. In contrast to all existing methods used to analyse the evolution of the mt gene order that only provide incomplete results, a complete logical approach will enumerate all the solutions and highlight the logical consequences.

In practice, calculating tree solutions is possible only for a small taxonomic dataset (model generation is NP-hard). But it is possible to analyse a broader taxonomic dataset by combining all the optimal solutions of smaller subdatasets (local analysis). In the present study, the tree solutions for the entire group of bilaterians group have been calculated thanks to the combination of 36 computations (see S3-S7 appendix). All the computations were done on a laptop with a 2.5 GHz processor.

## Acknowledgments

We wish to thank Claudine Chaouiya, Gabriel Nève and Daniel Papillon for helpful discussions and corrections of the manuscript.

## Author Contributions

Conceived and designed the study: LO and YP. Programming: LO. Mathematical proofs: PP. Wrote the paper: LO, PP, YP. All authors have seen and approved the manuscript

## Supporting Information

**S1 appendix.** Exact minimal distances matrix calculated with genome_comparison.c program.

**S2 appendix.** Logbook 1 - Chronological description of computations for Bilateria.

**S3 appendix.** Axioms and solutions for Deuterostomia.

**S4 appendix.** Axioms and solutions for Ecdysozoa.

**S5 appendix.** Axioms and solutions for Lophotrochozoa.

**S6 appendix.** Axioms and solutions for Bilateria.

**S7 appendix.** Axioms and solutions for Chaetognatha.

**S8 appendix.** Logbook 2 - Chronological description of computations for Bilateria - Annex for Chaetognatha.

**S9 appendix.** List of primary phylogenetic hypotheses (PPHs).

